# A Bayesian method to infer copy number clones from single-cell RNA and ATAC sequencing

**DOI:** 10.1101/2023.04.01.535197

**Authors:** Lucrezia Patruno, Salvatore Milite, Riccardo Bergamin, Nicola Calonaci, Alberto D’Onofrio, Fabio Anselmi, Marco Antoniotti, Alex Graudenzi, Giulio Caravagna

## Abstract

Single-cell RNA and ATAC sequencing technologies allow one to probe expression and chromatin accessibility states as a proxy for cellular phenotypes at the resolution of individual cells. A key challenge of cancer research is to consistently map such states on genetic clones, within an evolutionary framework. To this end we introduce CONGAS+, a Bayesian model to map single-cell RNA and ATAC profiles generated from independent or multimodal assays on the latent space of copy numbers clones. CONGAS+ can detect tumour subclones associated with aneuploidy by clustering cells with the same ploidy profile. The framework is implemented in a probabilistic language that can scale to analyse thousands of cells thanks to GPU deployment. Our tool exhibits robust performance on simulations and real data, highlighting the advantage of detecting aneuploidy from two distinct molecules as opposed to other single-molecule models, and also leveraging real multi-omic data. In the application to prostate cancer, lymphoma and basal cell carcinoma, CONGAS+ did retrieve complex subclonal architectures while providing a coherent mapping among ATAC and RNA, facilitating the study of genotype-phenotype mapping, and their relation to tumour aneuploidy.

**Author summary:** Aneuploidy is a condition caused by copy number alterations (CNAs), which brings cells to acquire or lose chromosomes. In the context of cancer progression and treatment response, aneuploidy is a key factor driving cancer clonal dynamics, and measuring CNAs from modern sequencing assays is therefore important. In this framing, we approach this problem from new single-cell assays that measure both chromatin accessibility and RNA transcripts. We model the relation between single-cell data and CNAs and, thanks to a sophisticated Bayesian model, we are capable of determining tumour clones from clusters of cells with the same copy numbers. Our model works when input cells are sequenced independently for both assays, or even when modern multi-omics protocols are used. By linking aneuploidy to gene expression and chromatin conformation, our new approach provides a novel way to map complex genotypes with phenotype-level information, one of the missing factors to understand the molecular basis of cancer heterogeneity.

## Introduction

Cancer is a disease in which cell subpopulations with enhanced functional capabilities emerge, evolve and undergo selection against the immune system response and treatment [1]. The investigation of tumour evolution in terms of omics layers – e.g., genome, transcriptome, proteome, epigenome, metabolome – has key translational repercussions [2], and can benefit from widespread single-cell sequencing technologies [3] that probe, among many, RNA (scRNA-seq) and ATAC (scATAC-seq) from biopsies and patient-derived model systems [4]. With the current technologies, the most recent protocols can also extract multiple measurements from the very same cell (e.g., G&T [5] or GoT [6] for matched RNA/ DNA, or 10x multiome [7] for ATAC/ RNA), even if “multimodal” technologies have still limited diffusion because they are very expensive and relatively low throughput. For this reason, a much more common single-cell design is based on separating cells before sequencing, with many computational efforts focused on integrating, a posteriori, data generated from different data modalities.

In this second scenario, sometimes referred to as diagonal integration [8], the general idea is to map the measurements in a latent space, using some unsupervised integration method. If we do not need the latent space to reflect any specific biological quantity, variational autoencoders or factor analysis can be adopted [9–12]. Otherwise, when it is required that the latents are biologically interpretable, other approaches should be preferred. In cancer genomics and tumour evolution studies, one possibility is to reconcile RNA and ATAC from the observation that both capture distinct aspects of the same DNA molecule. In this sense, RNAs are products of the transcriptional processes that initiate from DNA, and ATAC is an assessment of chromatin conformation, a physical feature of DNA. Therefore, an interesting attempt – which is the one we follow in this paper – is mapping RNA and ATAC on latent DNA states. Notably, an extra layer of complexity is introduced by observing that latent DNA configurations can differ between subgroups of cells characterised by multiple types of genetic mutations. Here we define these as tumour clones with associated Copy Number Alterations (CNAs), potent modifications of the chromosome conformation and copies.

While point mutations are difficult to characterise and link to RNA and ATAC, the opportunity of modelling latent CNAs seems more feasible. The possibility of inferring latent tumour subclones from scRNA-seq has been already widely investigated [13–17], and some preliminary attempts at working with scATAC-seq are also present [18–20]. In this framing, we recently introduced CONGAS [13], a method to perform CNA-based integration from scRNA-seq. Starting from a genome segmentation (set of breakpoints), CONGAS uses a Bayesian probabilistic model to infer latent total CNAs (i.e, per segments ploidy estimates) while clustering input cells. CONGAS was the first model to join signal deconvolution (i.e., clustering), while detecting subclonal patterns of aneuploidy, and worked better than methods like InferCNV [14], HoneyBADGER [21], CopyKAT [17] and Numbat [22] that decoupled CNA inference from clustering. The solution achieved with CONGAS was however only partially satisfactory, because the statistical signal of CNAs in scRNA-seq is generally affected by strong confounders such as allele-specific expression and post-transcriptional regulation, two biological phenomenon that are only partially understood and play an important role in cancer [23–25]. In practice, the distribution of read counts in RNA space (the inverse of the latent mapping), is not a perfect predictor of CNAs, and a better-quality signal can instead be achieved by examining chromatin conformation, a direct measurement of DNA. We reasoned that, as in CONGAS, one can postulate that the more alleles (i.e., copies) of a chromosome region are open, the stronger the signal of ATAC on the region should be. In this sense, a model a-la-CONGAS could be developed to link the latent CNA to the observable ATAC peaks. As far as we understand, such an intuition has never been leveraged before, missing the possibility of integrating independent scRNA-seq and scATAC-seq measurements using a biology-informed latent model of DNA copy numbers.

Building on this intuition, in this paper we present CONGAS+, a Bayesian model to map single-cell RNA and ATAC measurements in latent CNAs, clustering cells across the two data modalities, and predicting clones with a well-defined discrete copy number profile. Doing so, the CONGAS+ framework opens the possibility to compare both gene expression and chromatin accessibility across copy number clones. As a byproduct, the tool can readily separate tumour from normal cells when the former are characterised by aneuploidy, a very common situation in cancer. The model is unsupervised, and the likelihoods of RNA and ATAC are conditionally independent given the latent CNAs, but combined thanks to a shrinkage statistics to weight the contribution of the data modalities unevenly, which helps when one of the two modalities (usually RNA) is a worst predictor of CNAs. The overall model uses stochastic variational inference and gradient descent to learn parameters from data, and enjoys a fast implementation via probabilistic programming in Pyro [26]. This allows deploying CONGAS+ on GPUs seamlessly, analysing datasets with tens of thousands of cells in matters of minutes thanks to the massively parallel architectures offered by graphical devices. In this paper we show its capacity to extract complex genotype/ phenotype information from a B-cell lymphoma (∼ 6400 cells RNA/ATAC multimodal), a basal cell carcinoma (∼ 1200 cells RNA, ∼ 1200 cells ATAC) and a prostate cancer cell line (∼ 7600 cells RNA, ∼ 8800 cells ATAC).

## Materials and methods

### The CONGAS+ statistical model

CONGAS+ is a Bayesian model to infer copy number clones from scRNA-seq and scATAC-seq data, while simultaneously grouping cells into clusters characterised by similar Copy Number Alterations (CNAs). The model can also process data for which only one modality is available, as well as multimodal data generated by a multi-omics assay. Applied to cancer, CONGAS+ can i) separate tumours from normal cells (if the tumour has aneuploidy), ii) and detect tumour subclones associated with distinct CNAs. By default, the model scans for large CNAs at the resolution of chromosome arms but can be run also with a custom segmentation/ ploidy, for instance obtained by an independent sequencing assay. The model encodes this information as categorical priors on discrete copy number values, whose posteriors are obtained from a likelihood of scRNA and scATAC counts pooled at the resolution of the segments. The process of integrating two distinct measurements reflects also on input data resolutions: e.g., for ATAC we use either a fixed-width binning or open chromatin peaks, while for RNA we employ transcript counts. Notably, CONGAS+ is inspired by its predecessor CONGAS [13], the first method to attempt this inference from scRNA-seq alone. CONGAS+ has two main advantages: first, it builds a stronger statistical signal by joining ATAC on top of RNA, and second it models discrete CNAs, whereas the original model was approximating continuous copy numbers, making it difficult to compare the difference among clones in a solid statistical way.

CONGAS+ is a parametric Dirichlet mixture for *K* ≥ 1 clusters, determined from scRNA/ scATAC counts data *X* of *N >* 0 and *M >* 0 cells, respectively, in *I >* 0 segments. The model can process both discrete raw counts, as well as their continuous version normalised by library size and, interestingly, it can also work for a multimodal RNA/ATAC assay where *N* = *M*. As in other methods [13, 16], the total genome ploidy of a segment (sum of the latent major and minor alleles) is a linear predictor of the observed counts, i.e., if the latent number of DNA copies of a segment *i* is c, if we found *r >* 0 RNA transcripts and *a >* 0 open chromatin peaks mapped to the segment, then the predictors are of the form 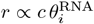 and 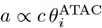 for certain parameterizations of the rate at which we observe RNA transcripts, and ATAC peaks.

The CONGAS+ likelihood of every modality (denoting its data broadly as *X*) is defined as

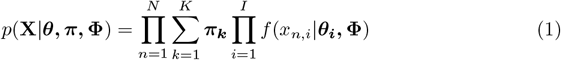

where *X* is a *N* × *I* matrix if there are *N* cells for the modality, *π* are the *K*-dimensional mixing proportions, *x*_*n,i*_ the counts for the *n*-th cell on the *i*-th segment, and Φ is a *K* × *I* × *H* tensor that models the probability distribution over latent CNAs, which here have *H* = {1, 2, … } possible states (usually capped at 5 copies). To map independent scRNA-seq/ scATAC-seq on the same set of clusters, Φ does not change across modalities. Note that *f* is a generic observational model that depends on counts either being discrete or normalised. The model is also available in two configurations, one with *π* independent across RNA/ATAC, and one where mixing is shared, which helps if signal strength is uneven across datasets.

For scRNA-seq/ scATAC-seq integer counts *x*_*n,i*_ mapped to the *i*-th segment of the *n*-th cell, the function *f* associated with the *k*-th mixture component is a Negative Binomial parameterized by mean and overdispersion, i.e.,

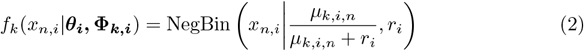

In CONGAS+ the mean depends on the expected counts per allele (*θ*_*i*_, learnt), the library size factor of the cell (*ρ*_*n*_, observed), and the linear combination (dot product) of the latent CNAs, i.e.,

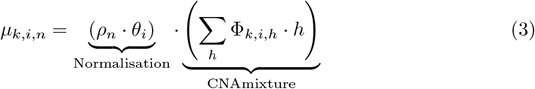

where we are using a nested mixture of CNAs with value *h* weighted by their probability Φ_*k,i,h*_ – i.e., the probability to detect CNA value *h*, in segment *i* and cluster *k*. The overdispersion of this segment is instead learnt from data. At the level of priors, the probability for each CNA Φ_*k,i,h*_ is a Dirichlet distribution with parameter an |*H*| -dimensional vector *α*, which can be set to any input predefined CNA profile: if we expect a ploidy *p* for a segment, we assign to the *p*-th entry value 0.6, and to the remaining 0.1 to obtain skewed samples from the Dirichlet. The joint scRNA/scATAC CONGAS+ likelihood for **X** = [*X*^*RNA*^, *X*^*AT AC*^] has a shrinkage form

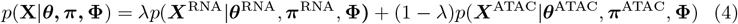

where 0 ≤ *λ* ≤ 1 is a fixed hyperparameter to weight the likelihood of both modalities.

For a given configuration with K clusters, parameters are learnt using a stochastic variational inference schema, giving in input segments that, from an isolated run, are fit to at least two clusters (i.e., multi-modal segments). Moreover, in learning the categorical Φ, we perform a reparameterization based on the Gumbel-softmax to draw low variance differentiable samples [27]. The inference returns the posterior distribution over copy number profiles and cell clustering assignments, while for all the other parameters CONGAS+ outputs the Maximum A Posteriori (MAP) estimate. The number of clusters *K >* 0 is optimised via model selection of standard score functions [28]: the Bayesian or Akaike Information Criteria (BIC, AIC), as well as the Integrated Completed Likelihood (ICL) [28]. In particular, given the complete log-likelihood *L*(**X**) = ln *p*(**X**|*θ, π*, Φ) and the number of parameters *v* for a model with *n* samples, the scores are BIC(**X**) = *v* ln (*n*) − 2*L*(**X**), AIC(**X**) = 2*v* − 2*L*(**X**) and ICL(**X**) = BIC(**X**) + *H*(*Z*) where *Z* are the latent variables for cell clustering assignments, and *H*(*Z*) their entropy [29]. To choose the optimal number of clusters we minimise any of those (BIC by default).

The full formulation of CONGAS+, including the alternative Gaussian continuous likelihood are provided as Supplementary Material. CONGAS+ is implemented in 2 open-source packages: one, in Python, using the probabilistic programming language Pyro [26], while the other, in R, to preprocess data and visualise inputs/ outputs, interacting with Python through reticulate.

## Results

### Model validation and parameterization

#### Performance and comparison to alternative methods

We performed simulations to assess the performance of CONGAS+ and other methods. To obtain an unbiased and realistic performance assessment, we simulated scRNA-seq and scATAC-seq data outside of CONGAS+, even if our method can generate data.

We simulated RNA and ATAC of normal cells via simATAC [30] and SPARsim [31], two tools for synthetic data generation calibrated on real sequencing technologies. Then, we added CNAs for 2 ≤ *K* ≤ 10 clones assembled from a random clone tree [32] with ≤ 5 extra CNAs every new subclone added to the tree. Finally, we generated mixing proportions from a Dirichlet distribution with uniform concentration, and assigned cells to clusters in 10 replicas for each K, for 90 total datasets, each one with 1500 scRNA-seq and 1500 scATAC-seq cells.

We applied CONGAS+ to each dataset searching for *K* ≤ 10, and measured i) the Adjusted Rand Index (ARI), i.e. the similarity between the known and inferred cluster memberships, and ii) the mean absolute error (MAE) between cluster-specific CNAs, and simulated ones. Across all tests we observed a very good performance (median ARI *>* 0.8 and MAE consistently lower than 1) suggesting that CONGAS+ can retrieve the cells from each clonal population, as well as their copy number profiles (Figure 1I,K). The same test has been carried out against alternative CNA detection algorithms that can work with either RNA or ATAC single-cell data (Figure 1J). In particular, we measured ARI for i) CONGAS+ ii) copyKAT [17], which uses only RNA data, and iii) copy-scAT [20], which uses only ATAC data. In this test we observed that CONGAS+ outperforms both tools. We explain this because our method can work even in the presence of pure tumour samples and multiple subclones, whereas the other two callers can only separate tumour from normal cells, therefore their usage is more limited.

**Fig 1.**
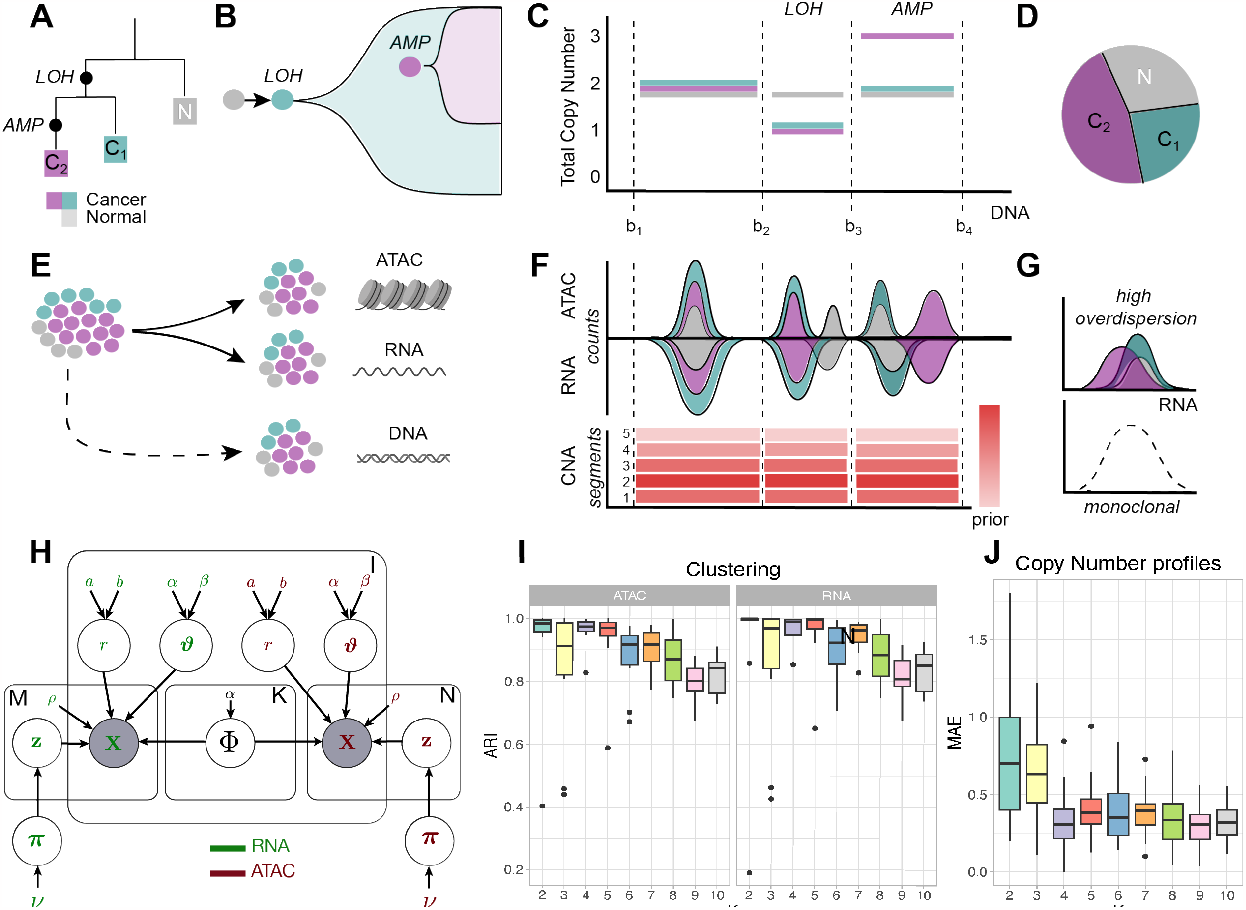
CONGAS+ framework. A-D: Given a tumour sample characterised by an accumulation of Copy Number events (A-B), CONGAS+ aims at inferring the total Copy Number profiles (C) and the clonal composition of each sample (D). E-F: CONGAS+ takes in input the single-cell RNA, single-cell ATAC and, optionally, a copy number segmentation obtained from a bulk whole genome assay, which is used as a prior for the copy number state of each segment. In case it is not provided, the tool employs a diploid arm-level segmentation. G: CONGAS+ infers the copy number state in each segment. H: Probabilistic Graphical Model. I: Adjusted Rand Index (ARI) computed comparing the ground truth labels of simulated cells with the clustering assignments returned by CONGAS+. 90 datasets with 1500 cells for scRNA and 1500 cells for scATAC were simulated. J: CONGAS+ performance compared with copyKAT [17] and copy-scAT [20], computing the ARI obtained on the same simulated datasets as in I. K: For the same data in panel (I), Mean Absolute Error (MAE) between the ground truth and the inferred copy number profiles. L: Boxplot showing the ARI for copyKAT, CONGAS and CONGAS+ (computed on scRNA and scATAC separately) obtained running the tools on bootstrap samples characterised by a bimodal signal poorly evident in RNA.

#### The importance of using a joint assay

CNA-associated signal quality is not necessarily even across ATAC and RNA, with the latter showing higher overdispersion due to difference in sample preparation, library size, gene expression variability, and sequence-specific biases [33, 34]. High overdispersion can act as a confounder, making it difficult to infer poorly separable clusters from scrNA-seq data alone, eventually leading to failures in detecting subclones. Using ∼ 1800 scRNA and ∼ 600 scATAC profiles from the Basal Cell Carcinoma (BCC) sample SU008 [35, 36], we created a dataset of tumour and normal cells in even proportions, subsetting the genome to two diploid and two with aneuploidy, with bimodal signal poorly evident in RNA. Then, we performed non-parametric bootstrapping for the genes in each segment, and compared 30 inferences with CONGAS+ (RNA plus ATAC), CONGAS (RNA) and copyKAT (RNA). Using a joint ATAC-RNA assay, CONGAS+ with *λ* = 0.1 detected CNAs that distinguish tumour from normal cells, obtaining a median ARI ∼ 0.7 on ATAC but a lower ARI on RNA (Figure 1L). In general, due to the weaker RNA signal, all tools that looked only at RNA struggled separating tumour and normal cells, with copyKAT and CONGAS unable to detect the split (Figure 1L). In this test, copy-scAT failed to execute with standard parameters. Overall, this shows that with a joint inference on the ATAC and RNA modalities we can detect the clonal structure of the dataset also when one data modality has a weak signal.

#### Shrinkage effect with Basal Cell Carcinoma data

Since signal quality can be uneven across data modalities, CONGAS+ is equipped with a shrinkage hyperparameter that can be used to weigh the evidence differently across RNA/ATAC. This serves as a natural hyperprior to decrease the importance given to a modality that we believe is more noisy or affected by some consistent bias. A natural question is therefore how does this affect the inference, and what value for *λ* should be suggested in the general case.

Building on the test shown in the previous section, we used data from SU008 and from sample SU006 [35, 36] in order to test a signal present in just ATAC (SU008), against a signal present in both RNA and ATAC (SU006). For SU006, we selected 2 diploid segments and 2 segments with monosomy loss of heterozygosity (LOH) (fig. 2F,G). To investigate the effect of *λ* on the inference, we scanned *λ* = 0.05, 0.15, …, 0.95, set *K* = 2 and performed 10 runs to compare the ARI for cluster assignments against tumour/normal labels from [35, 36]. We observed RNA/ATAC inferences stable against *λ*, with tumour and normal cells always separated (Figure 2C,H). For SU008, instead, only ATAC exhibits a neat bimodal distribution (fig. 2B), and this time we observed (fig. 2C) that for *λ <* 0.5 the ARI for ATAC is stable at ≈ 0.75, whereas it decreases as *λ* approaches 0.95. Inferences for the best/ worst ARI (fig. 2D-E) show discordant tumour and normal assignments. With *λ* ≥ 0.25 CONGAS+ did not fit the ATAC bimodality, merging 63% of tumour and 90% of normal cells together. Instead, for *λ <* 0.25 – more weight assigned to ATAC –assignments retrieved are perfect. As expected, the model is never able to separate tumour and normal from RNA, due to its unimodal distribution.

**Fig 2.**
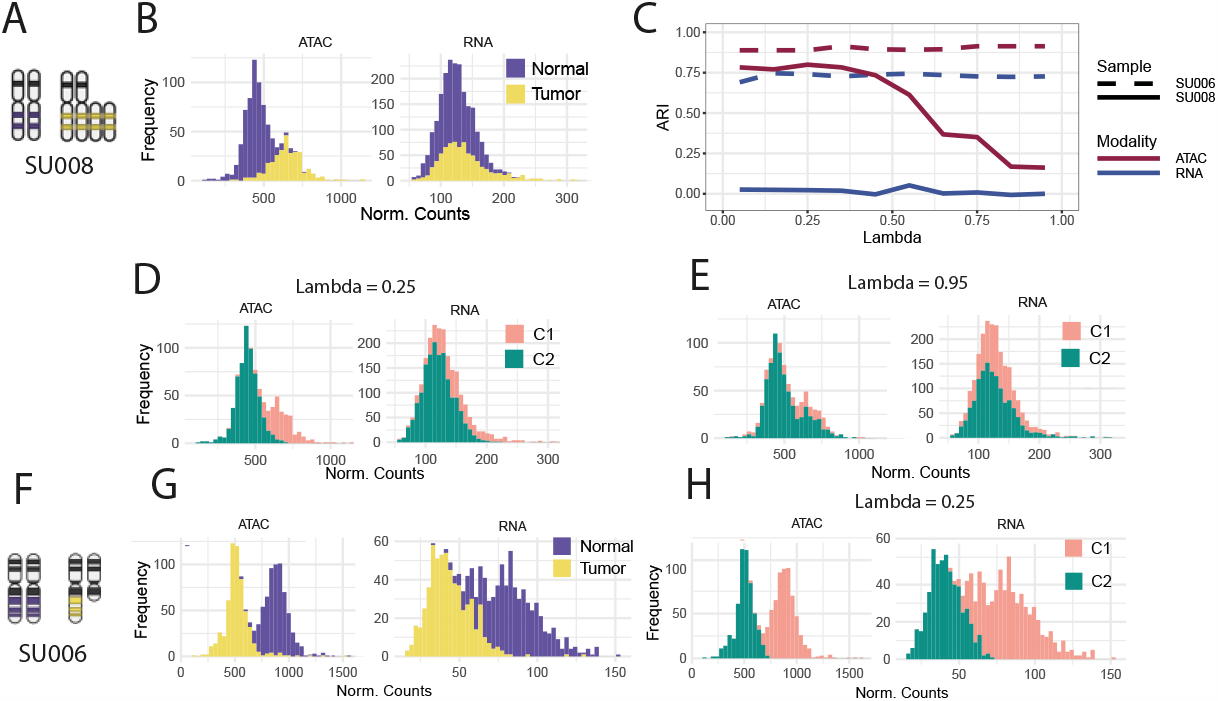
Impact of lambda variation on CONGAS+ performance using Basal Cell Carcinoma data. A,B,C,D,E: Basal Cell Carcinoma sample SU008 from [35] and [36], where we selected segments with bimodal signal in both scRNA-seq and scATAC-seq profiles. F,G,H: BCC sample SU006 from [35] and [36], where we selected segments in which ATAC signal is bimodal and RNA unimodal. C: ARI value for each modality, computed for every lambda ranging from 0.05 to 0.95. B,G: normalised counts distribution coloured by the ground truth cell labels for chromosomes chr9q and chr20q in samples SU008 and SU006 respectively. D,H: distributions coloured according to clustering assignments obtained from the solution showing the highest ARI for samples SU008 and SU006 respectively. E: normalised distribution for the worst solution in terms of ARI for sample SU008.

Overall, these tests show that if the quality of ATAC/ RNA are different, *λ* can be used to favour one assay over the other. CONGAS+ offers a principled approach based on likelihoods to inspect the optimal *λ*, and a final decision has to be taken on each dataset, also inspecting fits quality.

### Phasing ATAC and RNA profiles in B-cell lymphoma multimodal data

The ideal data to be integrated in CONGAS+ is a multimodal ATAC/RNA assay, where joint measurements are available for all cells. We gathered data of a B-cell lymphoma [37] sequenced with the 10x multiome kit [7], and tested if CONGAS+ did identify clusters across the two modalities, and assign cells consistently. In this test we phased cells across modalities to exploit the one-to-one correspondence between ATAC/RNA barcodes. The expectation was to cluster together cells in both RNA and ATAC, even without any a priori imputation. We processed ∼ 6400 cells with manually annotated (cfr. [37]) cell types after quality control. Cell types were distinguishable in a joint ATAC/RNA UMAP [38] low-dimensional representation (fig. 3A): two tumour cell populations (B and B-cycling) cluster together, whereas normal cells split into Monocytes, T and B cells. Note that, while CNAs could tell apart normal from tumour cells, the distinction among B and B-cycling tumour subpopulations is more likely linked to cell cycle entry dynamics, a byproduct of complex transcriptional regulation not necessarily linked to CNAs [39].

**Fig 3.**
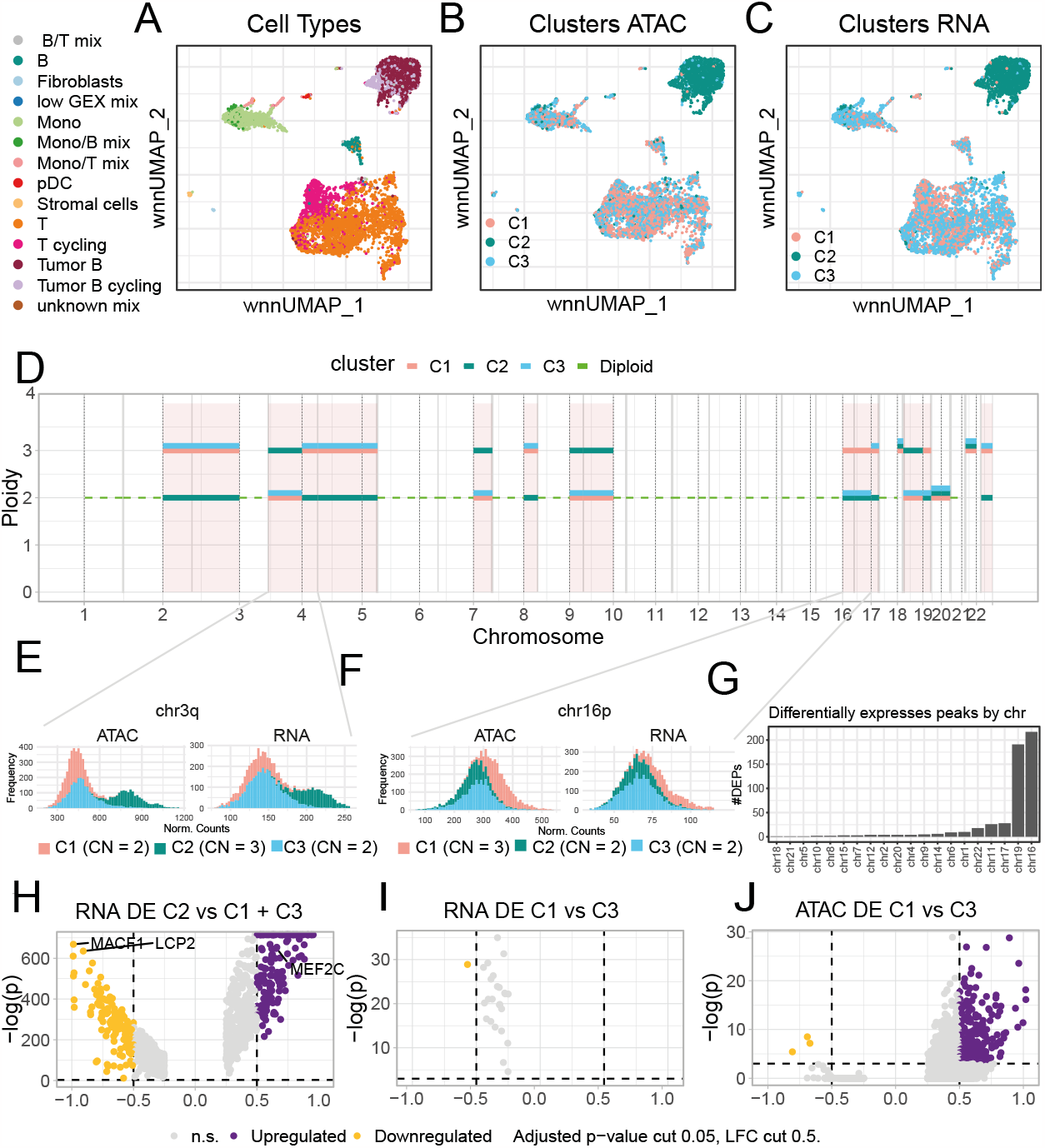
Application of CONGAS+ to B-cell lymphoma multimodal data. A: UMAP low-dimensionality representation of ∼ 6400 RNA and ATAC single-cell profiles from the 10x multiome dataset from [37]. B,C: The same UMAP plots coloured according to the clusters inferred by CONGAS+ for ATAC and RNA, respectively. D: copy number profiles for each cluster inferred by CONGAS+ E-F: normalised counts distribution coloured according to cluster assignments, shown for segment chr3q (E) where an amplification characterises tumour cells and chr16p (F) where an amplification is observed for cluster C1. G: distribution across chromosomes of the differentially expressed peaks between C1 and C3. H-I: volcano plot for differentially expressed genes between C1 and C3 (E) and between the tumour cluster C2 and normal cells. J: volcano plot for differentially expressed peaks between normal cells clusters C1 and C3.

We found *K* = 3 clusters using an arm-level segmentation and diploid priors (Figure 3D). Comparing cell labels from [37], we observed a single population of tumour cells, but two clusters composed of normal cells (Figure 3B-C). While the tumour/normal split is reasonably explained by CNAs which characterise neoplastic cells on chromosomes chr2, chr3q (Figure 3E), chr4, chr5p, chr7p, chr8p, chr9, chr18q and chr22q, the distinction among normals was unexpected and linked to distinct ATAC profiles for chromosomes chr 16p and chr 19 (Figure 3F). We begin to verify that CONGAS+ was not splitting normal cells due to differences in cell type composition. Then, we performed differential expression analysis across the RNA profiles of all populations. We did find – as expected – differences in the expression (Figure 3H) of genes that distinguish normal from lymphoma cells, as measured from absolute log fold change (LFC) above 0.5 and p-values below 0.01 (Wilcoxon test), but did not find differences across the two normal subpopulations (Figure 3I). Among the tumour-associated genes we find LCP2, a prognostic gene for metastatic melanoma-infiltrating CD8+ T cells [40], MEF2C a gene which has been linked with the lymphoma pathogenesis [41], and MACF1, a gene which has been associated with cancer development [42]. Instead, a differential analysis of ATAC peaks between the two normal subpopulations showed marked (|LFC| *>* 0.5 and p-values*<* 0.01; Wilcoxon test) ATAC differences across chromosomes chr16 and chr 19 (Figure 3G,J).

The possible explanation to this outcome could be that the subset of normals identified by CONGAS+, which cannot be otherwise identified by RNA-based tools, are an ancestor of the tumour population, describing the evolutionary link between normal cells and lymphoma cells. This will require further investigations also involving lineage-tracing via genetic polymorphisms to be confirmed, but this explanation would fit with biological observations that B-cell lymphomas originate from neoplastic transformation of germinal centre B cells [43]. Overall, this case study shows the combined power of ATAC and RNA, a joint framework that, to the best of our extent, has not yet been exploited before to study copy number alterations.

### CNA-associated drug-resistance clones in a prostate cancer cell line

Chromosomal instability and aneuploidy can generate potent phenotypes, sometimes capable of resisting negative selection induced by anticancer drugs [44]. We tested if CONGAS+ could identify, from scRNA-seq and scATAC-seq data, CNA-associated tumour subclones that resist treatment. We used data from [45], where ATAC/RNA data was generated for the untreated prostate cancer cell line LNCaP (parental), and then for one line treated 48 hours with AR antagonist enzalutamide (ENZ), and two resistant lines (RES-A and RES-B) derived after long-term exposure to ENZ and diarylthiohydantoin RD-162, respectively (fig. 4A). To search for high-resolution subclonal CNAs we downloaded LNCaP cytogenetics data from the DepMap portal [46], and used it to obtain breakpoint coordinates and priors for CNAs. We merged the 4 samples (parental, ENZ-48, RES-A, RES-B), and filtered out segments with more than 10% of cells showing zero counts.

**Fig 4.**
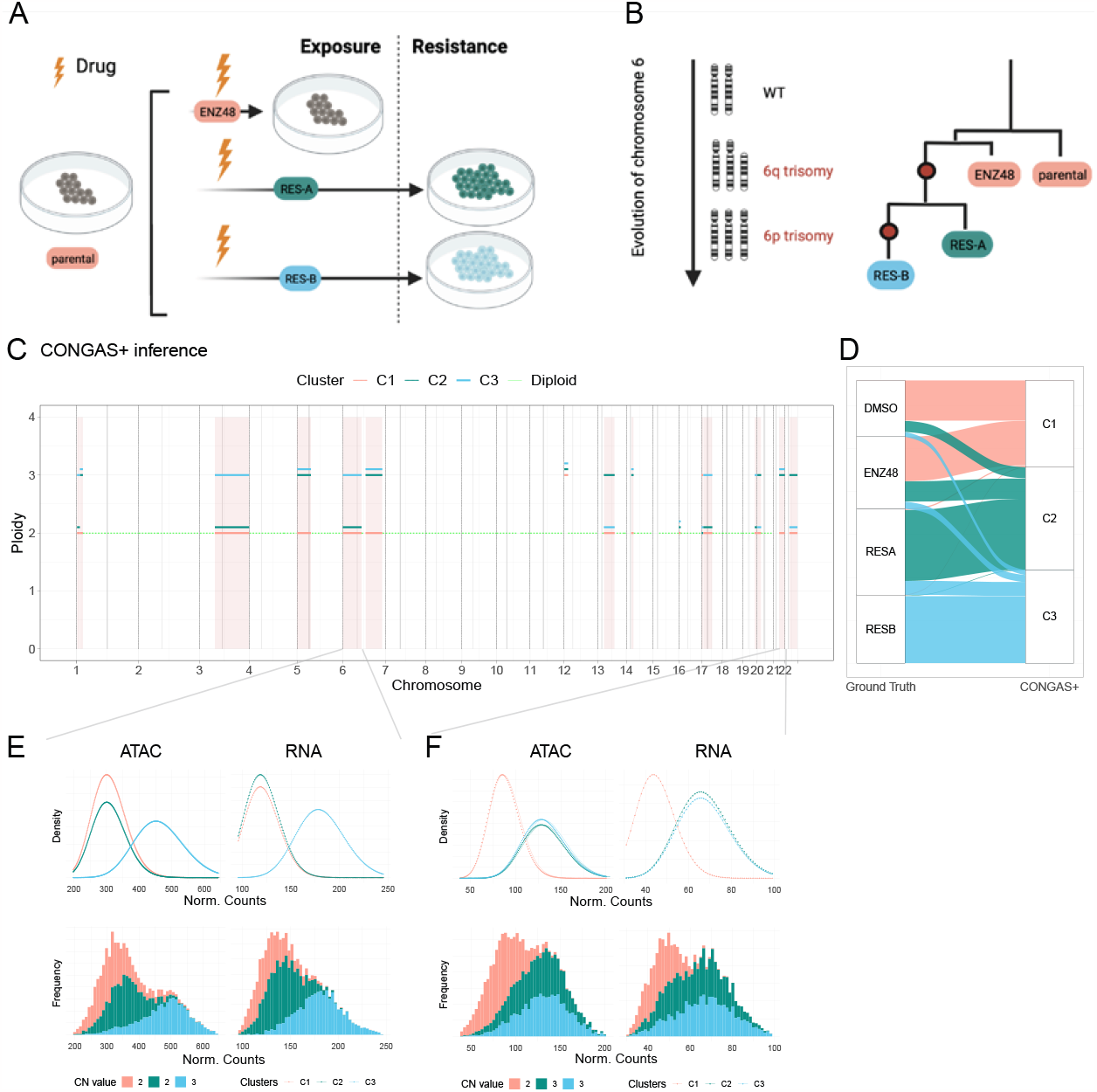
Application of CONGAS+ to prostate cancer cell line LNCaP with independent assays. CONGAS+ application to a prostate cancer dataset from [45], composed of a mixture of four cell lines with 7600 scRNA-seq cells and 8800 scATAC-seq cells. A-B: Cartoon representing the design of the drug resistance experiment and the corresponding sample tree. C: copy number profiles inferred by CONGAS+ for each cluster. D: Sankey plot showing the overlap between the sample of origin and the cluster inferred by the model. E-F: density plot and histogram of normalised counts coloured according to cluster assignments for chromosome 6p (D) where an amplification event is private to cluster C3, and chromosome chr21q where an amplification is shared by clusters C2 and C3.

CONGAS+ identified 3 clusters present in both ATAC and RNA across all samples (fig. 4C-F). The parental and ENZ-48 lines cluster together while RES-A and RES-B split in two clusters. This is consistent with the experimental design of [45]: ENZ48 has not yet acquired resistance due to its short-term exposure to ENZ, and is expected to cluster with parental cells. The two other clusters are composed of almost fully-resistant cells, with a good partition of the RES-A and RES-B cells. These two clusters share one amplification of the q-arm of chromosome 6 and 21 (fig. 4F), indicating evolution from sensitive cells through a common ancestor (fig. 4B). Moreover, the two resistant populations cluster separately and CONGAS+ finds, for RES-B, an amplification on the p-arm of chromosome 6 (fig. 4E), suggesting further evolution in that clone (fig. 4B).

Overall, this analysis shows that lineage relations associated with CNA-associated subclones can be effectively detected by CONGAS+ and longitudinal data, posing the bases for more systematic investigations on the causal roles of CNAs in promoting therapy resistance.

## Discussion

The relation between somatic mutations and cancer phenotypes is extremely complex and intimately related to the underlying evolutionary dynamics of cancer cells and the environment. To understand this genotype-phenotype mapping, single-cell technologies can be adopted to achieve a fine-grained resolution of the measurements, but methods are required to resolve signals in such noisy data. In this paper, we approached this problem from RNA and ATAC single-cell sequencing, inferring latent tumour subclones associated with CNAs, a specific type of complex genomic mutation. CONGAS+ is the first Bayesian model that can jointly analyse RNA and ATAC, inferring CNAs while clustering cells through variational inference. The model has a shrinkage formulation to weigh the evidence between the two modalities, a feature motivated by our experience where scATAC-seq has a cleaner signal while scRNA-seq is overdispersed. This phenomenon could be explained considering that ATAC is a direct measurement of DNA, while RNA is a byproduct and is therefore more subject to biases.

Using simulations, we assessed that CONGAS+ is robust and accurate in retrieving both the clonal composition and corresponding CNAs. This was further confirmed with real data, where the method showed the capacity to extract evolutionary relations that are difficult to retrieve with tools that analyse just RNA or ATAC. Moreover, we did appreciate the possibility of running CONGAS+ also on multimodal data, where RNA and ATAC are measured from the same cell. Even in this case, our model could correctly phase the sequenced cells across the modalities, suggesting that CONGAS+ is also ready to process multi-omics data.

In the future, following the stream of work on CONGAS and CONGAS+, we plan to further data modalities, e.g., methylations, which require ad hoc methods to be processed [47]. Moreover, another possibility is to include finer-resolution information such as B-Allelic Frequency (BAF) and Depth Ratio (DR) profiles, as commonly used to detect CNAs from bulk [48]. This would allow one to infer copy-neutral losses of heterozygosity which, otherwise, are diploid and therefore poorly identifiable. Moreover, BAF/DP profiles might also be exploited in conjunction with read counts data to implement an algorithm for de novo genome segmentation and copy number calling, without requiring any input bulk DNA data.

## Code and Data Availability

The CONGAS+ Python implementation and the R wrapping package are available at

- [Python] https://github.com/caravagnalab/CONGASp.
- [R] https://github.com/caravagnalab/rcongas/tree/categorical (*categorical* branch)

The code to reproduce the analyses in the text will be made available upon publication.

## Acknowledgments

This work was funded by AIRC under MFAG 2020 - ID. 24913 project – P.I. Caravagna Giulio, and by the CRUK/AIRC Accelerator Award #22790, “Single-cell Cancer Evolution in the Clinic” (MA, AG and GC).

