## Supplementary Material for "A Bayesian method to infer copy number clones from single-cell RNA and ATAC sequencing"

April 1, 2023

### 1 Additional Materials and Methods

#### 1.1 Relationship between single-cell signal and copy number.

The goal of CONGAS+ is that of identifying subsets of cells that are characterized by the same copy number value over each segment. We employ a bulk DNA sequencing experiment output to identify the segments, and we use their coordinates to aggregate the RNA and ATAC signal of each single cell over each segment. This choice is motivated by the goal of identifying a tradeoff between reliability of the model and overall cost of the analysis: compared to a single-cell assay, bulk DNA-seq experiments are characterized by a lower cost and they can be exploited to confidently identify Copy Number segments through a wide range of well-established tools, such as Sequenza [6], CNVkit [13] and GATK best practices [14]. Through this step we can model segments as independent entities, using as input to the model the aggregated counts for each single cell. CONGAS+ models the observed counts for each single-cell in each segment as variables dependent on the copy number state of that segment: once counts have been aggregated, we assume that the hidden copy number state influences the gene expression and chromatin accessibility signal through a linear relation: this is based on the intuition that if one tumor cell loses one of the two alleles in one segment, the genes on that segment will be characterized by a lower signal compared to the same genes in a normal (i.e., diploid) cell. This dependence has already been succesfully exploited on RNA in [5] and in our previous work [9], and we extended this to the chromatin accessibility signal: higher or lower Copy Number values for a specific segment predict more or less transcripts for those genes mapped to the segment, and amount of open chromatin on the segment.

#### 1.2 Full formulation of CONGAS+ statistical model

The model takes in input two single cell datasets $X^{atac}$ and $X^{rna}$ of sizes $N^{atac} \times I$ and $N^{rna} \times I$, containing respectively the signal of RNA and ATAC cells per segments. These two matrices are the result of a pre-processing step, were the counts are aggregated over each segment $i$ by summing all the features that map to $i$, with $i = 1, \dots I$. RNA features correspond to genes, while ATAC features correspond to peaks or fixed-length genome bins. Both these two matrices can either be non-normalised integer counts or normalised values, where in the latter case, we compute the z-score for each input feature (i.e., gene for scRNA and peak/genomic bin for scATAC). The type of input provided determines the distribution used to model the data, as it is explained in the next section.
CONGAS+ is a finite Dirichlet mixture of $K \geq 1$ distributions that model the $K$ clones present in the single-cell samples. The likelihood of our model has the following form:

$$p(\mathbf{X}^t | \boldsymbol{\theta^t}, \boldsymbol{\pi}, \boldsymbol{\Phi}) = \prod_{n=1}^{N^t} \sum_{k=1}^{K} \boldsymbol{\pi_k} \prod_{i=1}^{I} f(x^t_{n,i} | \boldsymbol{\theta^t_i}, \boldsymbol{\Phi}) \qquad (1)$$

where $N^t$ is the number of cells for modality $t$, $K$ the number of clusters and $I$ the number of segments. Here $f$ is a generic likelihood function which models the observed signal for the omic, $\pi$ are the clusters mixing proportions and $\Phi$ is a $K \times I \times H$ tensor the probability distribution over discrete copy number values for each cluster and each segment: each of the $k$ clusters is associated to a probability distribution per segment $\phi_{k,i,h} = P(C_{k,i} = h)$ over the possible copy number values $h = 1, .., H$ that the $i - th$ segment may assume, where by default, $H = 5$. The prior on the probabilities is a Dirichlet distribution

$$\Phi_{k,i,h} \sim Dirichlet(\alpha), \qquad \alpha = (\alpha_1, ...., \alpha_5), \ \ \alpha_i \in \mathbb{R}_{>0}, \tag{2}$$

where the concentration vector $\alpha$ is a hyperparameter chosen by the user. Given the ploidy $p$ of the segment, one may choose $\alpha = (\alpha_1 = 0.1, ...\alpha_p = 0.6, ...\alpha_5 = 0.1)$. Note that the tensor $\Phi$ does not change between modalities since cells from both omics are assigned to the same set of clusters.
CONGAS+ accommodates settings where the two omics have clusters in different proportions, which is achieved by using two different vectors $\pi^{atac}$ and $\pi^{rna}$, that model the mixing proportion for each cluster. Each entry $\pi_k$ is is sort by another Dirichlet distribution:

$$\pi_k^{rna} \sim Dirichlet(\nu^{rna}), \qquad \pi_k^{atac} \sim Dirichlet(\nu^{atac}) \tag{3}$$

where we choose $\nu^{rna} = \nu^{atac} = (1/K, ......, 1/K)$. Our model allows also to have a shared parameter $\pi$ to control the mixing proportions in ATAC and RNA jointly.

The generic likelihood function $f$ in Equation eq. (1) is defined based on the type of input matrix provided. On the one hand, for integer count matrices use Negative Binomial (NB) distributions to model the observed signal. On the other hand, with normalised counts each feature is z-scored prior to aggregating the signal over each segment and $f$ is defined as a Gaussian likelihood.

**Integer count matrix** With segment-specific integer counts we use the Negative Binomial (NB) likelihood,

$$p(x_{ni}^t | \rho_n^t, \theta_i^t, r_i^t, \Phi_{k,i}) = NB\left(\frac{\mu_{k,i,n}^t}{\mu_{k,i,n}^t + r_i^t}, r_i^t\right), \tag{4}$$

where $\mu_{k,i,n}^t$ and $r_i^t$ are the mean and size of the NB, respectively. The prior distribution on $r_i$ is $r_i \sim Unif(a_i^t, b_i^t)$ for some choice of the extrema $a_i^t, b_i^t$. The mean of the NB is defined as

$$\mu_{k,i,n}^t = \underbrace{(\rho_n^t \cdot \theta_i^t)}_{\text{Normalisation}} \cdot \underbrace{\left(\sum_h \Phi_{k,i,h} \cdot h\right)}_{\text{CNAmixture}} \tag{5}$$

Here $\theta_i^t$ are omic-specific variables that represent the average signal of a single copy of the $i$-th segment. For these quantities we choose a Gamma prior

$$\theta_i^t \sim \Gamma(\alpha_i^t, \beta_i^t), \tag{6}$$

where the hyperparameters $\alpha_i^t, \beta_i^t$ can be estimated from the data.
Through the definition of the Negative Binomial mean described in eq. (5), we are modeling our assumption that the observed signal for each cluster in each segment is proportional to the number of DNA copies of that segment for that specific cluster. In fact, the distribution $\Phi_{k,i}$ models the probability to detect each CNA value $h \in 1, \ldots, H$ (by default $H = 5$) for each cluster $k$ and segment $i$, and thus the mean of the NB depends on the linear combination (dot product) of the latent CNAs.
We also introduce the cell specific normalization factors $\rho_n^t$, which take into account possible expression differences due to sequencing. These are hyperparameters of the model and can be estimated from the data.

**Normalized count matrices** CONGAS+ supports also datasets where where prior to aggregation, each feature has been z-scored. In this case we assume the aggregated signal over each segment to be normally distributed, with the mean equal to the copy number value:

$$p(x_{ni}^t | \Phi_{k,i}, \sigma_i^t) = N\left(\mu_{k,i}, \sigma_i^t\right) \tag{7}$$

where

$$\mu_{k,i} = \sum_h \phi_{k,i,h} \cdot h \tag{8}$$

and the standard deviation $\sigma_i$ is sorted by a uniform distribution

$$\sigma_i \sim Unif(a_i^{rna}, b_i^{rna}). \tag{9}$$

The definition of mean of the Gaussian presented in eq. (8) is analogous to that of the Negative Binomial in eq. (5): $\Phi_{k,i}$ is the distribution over possible discrete CNA values $h \in 1, \ldots, H$ (by default $H = 5$) for each cluster $k$ and segment $i$, and thus the mean of the Gaussian depends on the linear combination of the latent CNAs.

**Copy number recalibration** CONGAS+ has one optional parameter that can be exploited to correct the copy number calls according to the input provided with the bulk. In detail, for each omic $t$ and each segment $i$ where $p_i$ bulk ploidy input, we compute the following penalization factor $s_i^t$:

$$s_i^t = \left( \sum_k \pi_k^t \cdot \sum_h \phi_{k,i,h} \cdot h \right) - (p_i \cdot \eta + 2 \cdot (1 - \eta)) \tag{10}$$

Where $\eta$ is the tumor purity of the sample, which can be set using the value estimated from bulk DNA. This factor $s_i^t$ serves as a penalization for solutions that deviate too much from the input bulk ploidy. The total penalization is computed as $s = \sqrt{\sum_i (s_i^t)^2}$, and it is subtracted from the total likelihood $p(X^t|\Delta^t, \Phi, \pi^t)$.

**Parameters estimates** Our inference algorithm requires marginalizing the likelihood with respect to the clustering and copy number assignment and calculating the Maximum A Posteriori (MAP) MAP estimates of the continuous parameters.
Once the MAP estimators have been computed, we can compute the copy number profile of each cluster $C_{k,i}$ by taking $C_{k,i} = \arg\max_h (\phi_{k,i,h})$. Given also the copy number states, one can compute the clustering assignment probabilities $P_{n,k}^t$ of the cells for both modalities. These are

$$P_{n,k}^t = \frac{\pi_k^t \prod_i f\left(x_{n,i}^t|\Phi_{k,i}, \theta^t, \rho^t\right)}{\sum_k \pi_k^t \prod_i \left(f\left(x_{n,i}^t|\Phi_{k,i}, \theta^t, \rho^t\right)\right)} \tag{11}$$

Using the above probabilities one can estimate for each cell the assignment vector $z_{n,k}^t$

$$z_{n,k}^t = \begin{cases} 1 \text{ if } k = \text{argmax}\left(P_{n,k}^t\right) \\ 0 \text{ otherwise} \end{cases} \tag{12}$$

##### 1.3 Variational inference for parameter estimation

Given the model definition presented above, we want to estimate the values for all parameters by learning their posterior distribution, defined using the Bayes rule. For simplicity, we use $U$ to indicate all the parameters in the model and thus we can write the posterior as

$$P(U|X) = \frac{P(X|U)P(U)}{P(X)} \tag{13}$$

Where $P(X)$ is the marginal likelihood, also called evidence. The denominator is usually intractable, and we need to approximate the real posterior. CONGAS+, like CONGAS uses Stochastic Variational Inference (SVI) [11] to get the approximation of the true posterior $p(U|X)$. The aim of SVI is to find a variational distribution $q(U)$ that belongs to a family of probability distributions $\mathcal{Q}$ and can approximate the real posterior. This can be formulated as an optimization problem, where the goal is to minimize the Kullback-Leibler (KL) divergence between $p(U|X)$ and $q(U)$ [11, 4]:

$$q^*(U) = \underset{q(U)\in\mathcal{Q}}{\arg\min}\{KL(q(U)||p(U|X))\} \tag{14}$$

However, this term is still untractable, as it requires to compute the posterior. Thus, the objective function that gets optimized in SVI is the Evidence Lower Bound (ELBO):

$$\text{ELBO}(q) = \mathbb{E}\left[\log p(U, X)\right] - \mathbb{E}\left[\log q(U)\right]. \tag{15}$$

and maximizing this quantity is equivalent to minimizing the KL divergence [8, 3, 11]. In detail, the variational distribution $q$ is parametrized by $\gamma$, which is what we want to learn during the inference, and in order to optimize the ELBO, SVI computes gradient descent optimization taking a Monte Carlo estimates of the gradient, which is defined as:

$$\nabla\gamma\text{ELBO} = \nabla_{\gamma}\mathbb{E}_{q_{\gamma}(U)}\left[\log p(x, u) - \log q_{\gamma}(U)\right]. \tag{16}$$

CONGAS+ is implemented in `Pyro` [2], a probabilistic programming language based on `Python` which implements SVI, we use *Adam* as an optimizer.

#### 1.4 The Gumbel-Softmax distribution

CONGAS+ uses SVI to approximate the posterior by performing gradient descent. However, our model contains a discrete random variable $\Phi$, that encodes the probability distribution over the possible discrete copy number values. In order to be able to estimate the gradient for this categorical variable, we use the Gumbel-softmax [7] (GSM), which is a continuous distribution that can approximate samples from a categorical distribution.
Considering the categorical probability distribution $\Phi$ which has $H$ probability classes $\alpha_1, \alpha_2, \ldots, \alpha_H$ and models the distribution over the possible copy number values, a sample $\omega$ from such distribution can be seen as a one-hot vector that lies on the corners of a $(h-1)$-dimensional simplex $\Delta^{h-1}$:

$$\omega = \mathit{one\_hot}(\text{argmax}_h[g_h + \mathit{log}\alpha_h]) \tag{17}$$

where $g_h$ are i.i.d. samples drawn from Gumbel$(0, 1)$ [7].

In order to approximate the argmax to make it continuous and differentiable, the softmax is employed. Thus, a $H$-dimensional sample vector from the Gumbel-softmax distribution is a vector $\omega \in \Delta^{h-1}$ defined as:

$$\omega_h = \frac{\exp\left((\mathit{log}(\alpha_h) + g_h)/\tau\right)}{\sum_{j=1}^{k}\exp\left((\mathit{log}(\alpha_j) + g_j)/\tau\right)}$$

Where $i = 1, 2, \ldots k$ and $\tau$ is the temperature parameter. As $\tau$ approaches 0, the samples from the Gumbel Softmax become one-hot vectors and thus sampling from the GSM becomes identical to drawing samples from the categorical distribution $\Phi$.

In the GSM definition, $\alpha$ is the vector of parameters of the distribution, and it corresponds to the vector of the probabilities for the $H$ classes of the categorical distribution. For values of the temperature greater than zero, the GSM has a well defined gradient with respect to its parameters, and thus if we replace the categorical samples with the Gumbel Softmax it is possible to use backpropagation during training to compute the gradients.

However, the samples from the GSM are not identical to samples from the corresponding categorical distribution when $\tau$ is not zero and thus there is the need for identifying a trade-off between large and small temperatures. In fact, on the one hand for temperatures close to zero samples are close to one-hot, but the variance of the gradients is large. On the other hand, large temperatures yield small gradient variance but smooth samples. The solution is to decrease the temperature followig a schedule: in Pyro we start from a value $\tau_{start}$, and then at each step $j$ of gradient descent optimization we use the temperature $\tau_j = \tau_{start}/\log(j + 0.1)$.

### 2 Additional Results

#### 2.1 Additional details on simulations

##### 2.1.1 Normal population simulation

In order to perform extensive simulations of CONGAS+, we have generated synthetic datasets. To obtain coupled scRNA and scATAC datasets, we employed two tools that simulate data by estimating parameters from real single-cell datasets, namely SPARSim [1] and simATAC [10]. In detail, we downloaded two public datasets of human Peripheral Blood Mononuclear cells (PBMC) cells from 10x genomics, and after performing quality check and clustering through Seurat and Signac we associated a PBMC cell type to each cluster with the tool MAESTRO, which exploits known gene expression signatures to assign labels based on cluster markers. We selected monocytes and neutrophils from scRNA and scATAC respectively, and we used their cells to obtain simulation parameters and generate synthetic datasets via SPARsim and simATAC.

simATAC [10] takes in input a bin x cells matrix and uses the real values to estimate the distribution of following simulation parameters: library size, non-zero cell proportion in each bin and mean signal for each bin. After sampling the library size for each cell i and the bin mean for each bin j, it simulates counts cij sampling from a Poisson distribution, where the mean of the Poisson is the library size scaled by the bin mean. The final output of the simulation is a bin x cells matrix of an homogeneous population of cells.

SPARsim [1] takes in input a single cell gene expression matrix and exploits it to estimate the model parameters, which are the following: feature intensity (i.e., vector representing the average expression level of each gene), biological feature variability and library size.

##### 2.1.2 Copy number clones simulation

We then added a copy number to the simulated normal data, considering simulated scenarios with increasing complexity in terms of clone-composition.

To simulate segment breakpoints we follow this procedure: we first divide the length of each chromosome $c$ by a parameter that encodes the approximate segment length (by default we set it to 1e8), to obtain the maximum number of segments $imax_{chr_c}$. Then, we randomly sample an integer $i_{chr_c}$ ranging from 1 to $imax_{chr_c}$, which corresponds to the effective number of segments. Finally, we break each chromosome in exactly $i_chrc$ segments and we randomise the breakpoint by adding a value sampled from a Uniform distribution $Unif(-a, a)$ to the segments coordinates, where $a$ is equal to 10% of the average segment length.

In order to simulate a clonal architecture, we generate a tree with $K$ clones by iteratively attaching a new node to a randomly selected leaf in the tree. The aneuploidy profile of each clone $y_k = (y_{k,1}, \dots, y_{k,I})$ is simulated by taking the parent profile and changing the copy number values of $D$ randomly selected segments. Through this procedure, each new node will have a Hamming distance from its parent equal to $D$. Once the clonal architecture has been computed, we divide the normal simulated cells in $K$ clusters, sampling their proportions from a Dirichlet distribution $Dirichlet(1/K, \dots, 1/K)$, and for each segment $i$ and each cluster $k$ we take all the features mapping to $i$ and we multiply their value by $\frac{y_{k,i}}{2}$.

We simulated datasets with $K$ ranging from 2 to 10, for a total of 90 datasets.

#### 2.2 Additional details on real-world datasets

##### 2.2.1 Basal Cell Carcinoma

We collected data from [15] and [12] where the authors performed scRNA-seq, Whole Exome sequencing (WXS) and scATAC-seq on samples collected from patients affected by Basal Cell Carcinoma (BCC). Patients underwent treatment against BCC, and sequencing experiments were performed both pre and post treatment. The datasets consist of both tumor and normal cells, and in order to assess the ability of CONGAS+ in identifying Copy Number clusters with varying values of $\lambda$, we considered the single-cell labeling provided by the authors, and we tested our framework on those datasets for which there was a significant presence of both tumor and normal cells. First, we analyzed patients SU006 and SU008 where all three assays were performed on both tumor and normal tissues, in order to assess the ability of CONGAS+ to identify Copy Number events that distinguish tumor from healthy diploid cells. We downloaded processed single-cell data and cell annotations from [15] and [12], and we selected an equal number of tumor and normal cells to build a dataset for illustrating the performance for varying values of hyperparameter $\lambda$. For bulk segmentation, we downloaded FASTQ files of both tumor and normal cells from SRA and we applied GATK [14] best practices for alignment: we performed alignment with `bwa-mem`, we removed duplicates with Picard and we recalibrated base quality scores. Finally, we used the obtained bam files as input for Sequenza [6] to identify CNA profiles.
